# Structures redefine the mechanism of action for tetracyclines

**DOI:** 10.1101/2025.09.25.678686

**Authors:** Swapnil C. Devarkar, Ivan B. Lomakin, Jimin Wang, Ayman Grada, Christopher G. Bunick

**Affiliations:** Department of Molecular Biophysics and Biochemistry, Yale University, New Haven CT, 06511, USA; Department of Dermatology, Yale University School of Medicine, New Haven, CT 06520, USA; Department of Dermatology, Case Western Reserve University School of Medicine, Cleveland, OH, 44106, USA; Program in Translational Biomedicine, Yale University School of Medicine, New Haven, CT 06520, USA

**Author notes:** These authors contributed equally to this work (Swapnil C. Devarkar, Ivan B. Lomakin).

## Abstract

The tetracycline class of antibiotics is widely used for treating bacterial diseases including Lyme disease, anthrax, acne vulgaris, and pneumonia. Using a series of high-resolution cryo-electron microscopy (cryo-EM) structures, we show that tetracyclines can simultaneously target the mRNA decoding center and the nascent peptide exit tunnel (NPET) of the bacterial 70S ribosome. Among the tested tetracyclines, Doxycycline was unique in its ability to dimerize and bind the NPET at multiple locations. Structural comparison of Doxycycline, Minocycline, and Sarecycline bound to the *Escherichia coli* and *Cutibacterium acnes* 70S ribosome revealed species-specific differences affecting drug interaction and occupancy. Our results redefine the mechanism of action for tetracyclines and provide a structural basis for rational design of narrow spectrum tetracyclines to overcome the rising threat of antibiotic resistance.

## Introduction

Originally discovered in the 1940s, the tetracycline class of antibiotics has been widely prescribed over the past seven decades for combating gram-positive, gram-negative, and atypical bacterial infections, and remains the preferred mode of treatment for diseases like acne vulgaris, respiratory tract infections, and tick-borne illnesses *(1)*. The tetracycline class of antibiotics derives its name from the four-benzene (naphthacene) ring core conserved across all generations (Fig. 1A). Tetracycline, initially approved by the U.S. Food and Drug Administration (FDA) in 1953, remains in production to date but its use has declined over time due to the rise of antimicrobial resistance (AMR). The second-generation tetracyclines, Minocycline*(2)* and Doxycycline*(3)*, were FDA-approved in the 1970s and carry substitutions and additional modifications to the naphthacene core at the C5, C6, and C7 positions that have led to enhanced efficacy and bypassed some of the evolved AMR mechanisms against tetracycline. The recently approved third-generation tetracyclines like Sarecycline*(4)* and Eravacycline*(5)* carry bulkier modifications at the C7 and C9 positions, respectively.

**Figure 1.**
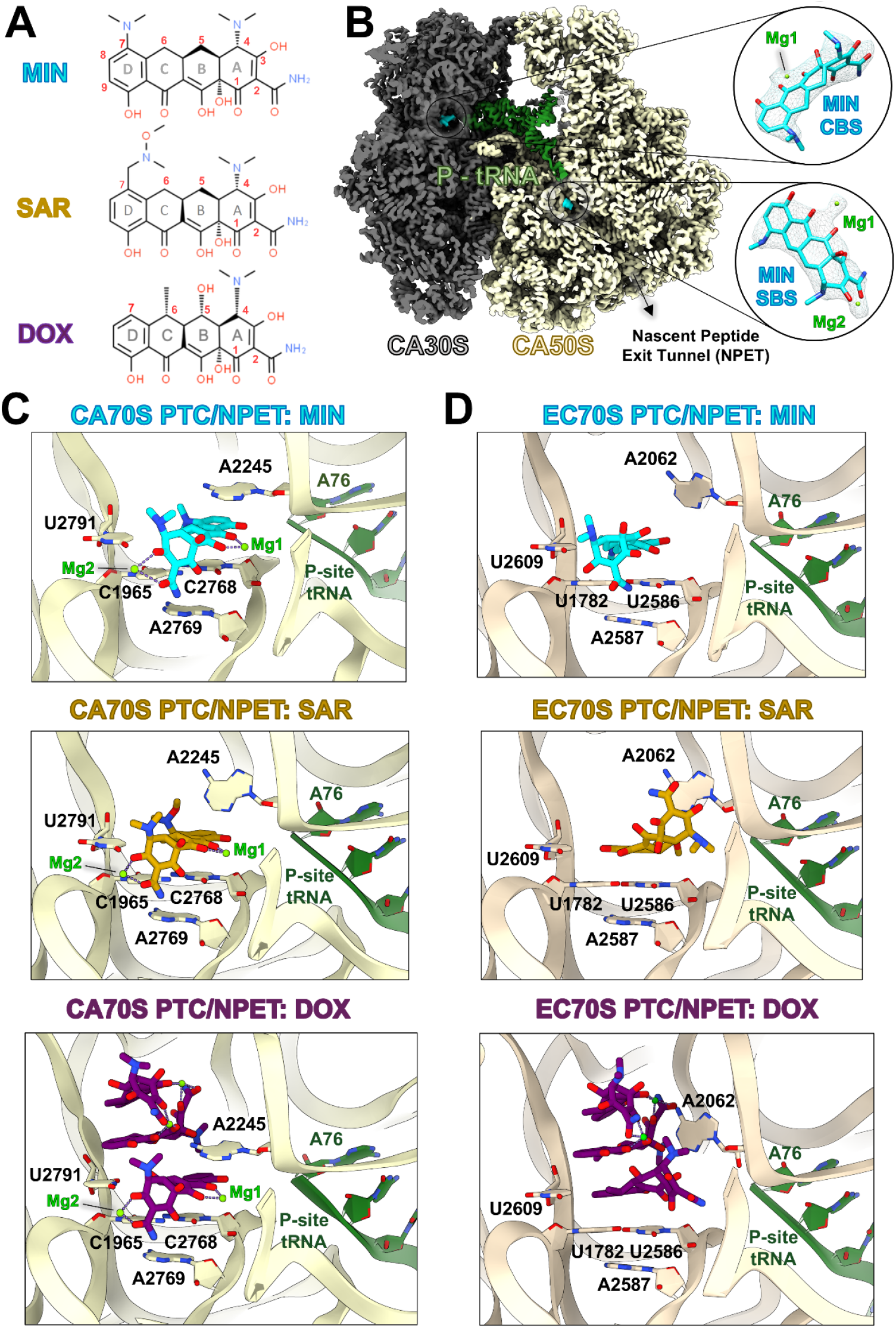
Tetracyclines bind the NPET of bacterial 70S ribosome. **(A)** The chemical structures of the three tetracyclines used in this study, Minocycline (MIN), Sarecycline (SAR), and Doxycycline (DOX) are shown in a schematic format. **(B)** Cryo-EM reconstruction of *C. acnes* 70S (CA70S) ribosome bound to Minocycline is shown. The insets highlight the atomic model, and the corresponding cryo-EM density of Minocycline bound at the canonical binding site (CBS) and secondary binding site (SBS). **(C-D)** Detailed views of the SBS of Minocycline, Sarecycline, and Doxycycline in the CA70S (C) and the *E. coli* 70S ribosome (EC70S) (D) are shown.

Current knowledge on the mechanism of action for tetracyclines states that tetracyclines abrogate bacterial protein translation by binding to the bacterial ribosome. Structural studies identified the mRNA decoding center in the 30S ribosomal subunit as the primary binding site, wherein it binds close to the acceptor site (A-site) codon and competes with the incoming A-site tRNA *(6-9)*. Some structural studies reported secondary binding sites for tetracyclines, but these sites were deemed inconsequential due to their distance from the two active centers of the bacterial ribosome, the mRNA decoding center and the peptidyl-transferase center (PTC), or considered an artifact of high antibiotic concentration *(6, 9, 10)*. Recently, we showed that Sarecycline binds at both active centers of the *Cutibacterium acnes (C. acnes)* 70S ribosome, at the canonical binding site (CBS) in the mRNA decoding center and at a secondary binding site (SBS) in the NPET near the PTC *(11)*. Recent structural studies on Tetracenomycin X, an aromatic polyketide with a four-ring structure similar to tetracyclines, demonstrated it targets the NPET of bacterial ribosomes and shares its binding site with the SBS of Sarecycline *(12, 13)*. However, unlike Sarecycline, Tetracenomycin X does not bind the 30S subunit of the bacterial ribosome and its inhibition of bacterial protein translation is mediated completely through its binding in the NPET. Whether other tetracyclines can target the NPET of bacterial ribosomes, similar to Sarecycline, remains uncharacterized.

A key advantage of tetracyclines is their versatility in mode of administration. The most common route is oral formulations but depending on the type of infection or disease being treated, topical, intramuscular and intravenous routes are also used. The effective concentration of tetracycline at the site of infection can vary widely depending on the dose and mode of administration *(14)*. For example, a doxycycline oral dose of 200 mg per 6 hours (typical dose ranges between 100-500 mg per 6 hours) leads to a peak plasma concentration of ∼7mg/l (∼15 µM) and ∼300 mg/l (∼700 µM) in the urine of healthy individuals *(15-17)*. Since Doxycycline is cleared from the body unchanged, it is used to treat gastrointestinal and urinary tract infections *(15)*. On the higher end of the spectrum, Minocycline (4%) topical foam, FDA-approved for treatment of moderate to severe acne vulgaris *(18)*, has a local concentration of Minocycline in excess of 80 millimolar. Whether the concentration of tetracyclines at their effector site impacts their mechanism of action remains unknown. Our current knowledge of tetracyclines stems from structural studies on model bacteria like *Escherichia coli (E. coli)* and *Thermus thermophilus (T. thermophilus)*. However, recent studies indicate that species specific differences in the architecture of the 70S ribosomes from pathogenic bacteria like *C. acnes* can influence the antibiotic binding and mechanism of action *(11, 19)*. Therefore, structural studies of tetracyclines bound to their clinical targets are needed to understand their mechanism of action and relevance to patient care.

To gain a deeper understanding of the mechanism of action for tetracyclines and the effects of species-specific differences in the 70S ribosome, we determined high-resolution cryo-EM structures of Minocycline, Doxycycline, and Sarecycline with the 70S ribosomes from the well-characterized gram-negative bacterium *E. coli* as well as the acne-causing gram-positive anaerobic bacterium *C. acnes*. Surprisingly, all the tetracyclines tested exhibited a two-site binding mechanism, the CBS in the 30S subunit and a SBS in the NPET adjoining the PTC. Doxycycline was unique in its ability to dimerize in the NPET, and two dimer pairs were observed in the NPET of both the *E. coli 70S* and *C. acnes* 70S ribosome. Interestingly, species-specific ribosomal differences lead to altered interaction networks and binding poses for the tetracyclines. The SBS of tetracyclines showed poor occupancy at lower concentrations, indicating that at higher concentrations (∼ 200 µM and above) bacterial translation is inhibited at the mRNA decoding center as well as the PTC. The catalog of high-resolution structures of three different clinically used tetracyclines bound to the enteric, gram-negative *E. coli* and the acne-causing, gram-positive *C. acnes* 70S ribosome provide a structural framework for leveraging the species-specific differences in the bacterial 70S ribosome in the rational design and development of future narrow-spectrum tetracycline derivatives.

## Results

### Tetracyclines bind the NPET of bacterial 70S ribosome

Prior structural studies on tetracycline established it binds to the bacterial 30S ribosomal subunit in the mRNA decoding center, blocking the A-site tRNA and inhibiting bacterial protein translation *(7, 8)*. Our structural investigation of Sarecycline, a third-generation tetracycline, with the 70S ribosome of the model bacterium *Thermus thermophilus* also showed binding to the same site in the 30S ribosomal subunit *(19)*. However, we recently reported that Sarecycline exhibited a two-site binding mechanism against *C. acnes* 70S ribosome, simultaneously targeting the mRNA decoding center in the 30S subunit and the NPET in the 50S subunit *(11)*. To gain structural insights into translation inhibition by other clinically prescribed tetracyclines, we determined high-resolution cryo-EM structures of the *C. acnes* and *E. coli* 70S ribosome in complex with Minocycline, Doxycycline, and Sarecycline (Figs. 1, S1, and Tables S1-S2). Surprisingly, in contrast to the current understanding on tetracyclines, Minocycline and Doxycycline also simultaneously targeted the *C. acnes* and *E. coli* mRNA decoding center and NPET (Figs. 1B-D and S2). All three tested tetracycline derivatives showed conserved structural features in their interaction network with the *C. acnes* NPET. The mismatch base pair (bp) C1965:C2768 in the *C. acnes* NPET plays a key role in mediating binding to the tetracyclines, wherein the rings B and C of tetracycline stack against C1965:C2768 bp at a perpendicular angle. Ring D is proximal to the peptidyl-transferase center (PTC) whereas ring A points towards the NPET (Fig. 1C). A2245 and U2791 provide additional stacking interactions for this secondary binding site (SBS) for tetracyclines. The conformation of the U2791 base in the Doxycycline bound *C. acnes* 70S structure is different compared to the Minocycline and Sarecycline bound ones due to the Doxycycline specific C5-hydroxyl forming hydrogen bonds with the N3 group of U2791 (Fig. 1C and S2A). All three tetracyclines coordinate two magnesium ions in the *C. acnes* NPET binding site. The first magnesium ion is coordinated by the hydroxyl atoms of rings B and C along with the phosphate backbone of C2623. The second magnesium ion is coordinated by ring A via its C3-hydroxyl and C2-carboxyamide group. Compared to the structure of Sarecycline bound to the *C. acnes* 70S ribosome reported herein, in our previous study *(11)*, the binding pose of Sarecycline in the SBS was modeled with a 180° flip. The higher resolution and cryo-EM map quality of the reconstruction reported herein allowed us to model the pose of Sarecycline in the SBS more accurately (Fig. S2B). The binding pose and interaction network of Minocycline, Doxycycline, and Sarecycline at the SBS in the NPET of the *C. acnes* 70S ribosome is highly conserved.

Minocycline, Doxycycline, and Sarecycline bind the *E. coli* NPET in the same location as in the *C. acnes* NPET. From here on all *E. coli* rRNA numbering is denoted in italics to differentiate from *C. acnes* rRNA numbering. In *E. coli* NPET, *U1782:U2586* forms the mismatch base pair, stabilized by *A2587*, that all three tetracyclines stack on (Fig. 1D and S2C-D). This stacking interaction is further stabilized by *A2062* and *U2609*. However, in contrast to the *C. acnes* NPET, the binding of the tetracyclines in the *E. coli* NPET showed greater variability. Minocycline and Doxycycline bind the *E. coli* NPET in a similar conformation to the *C. acnes* NPET but exhibited a rotation along the horizontal axis. Doxycycline exhibited a more dramatic change than Minocycline, with a ∼90° rotation (Fig. S3A). Sarecycline, on the other hand, binds the *E. coli* NPET in a different conformation to that observed for *C. acnes* NPET (Fig. 1D and S2D). Sarecycline is flipped by ∼180° and exhibits a similar axial rotation to Doxycycline. A probable reason for the flipped conformation of Sarecycline is its bulky C7 moiety which combined with the axial rotation would lead to a steric clash with *U2609*. The flipped conformation allows the C7 moiety to be accommodated in the pocket. In contrast to the SBS in *C. acnes* NPET, none of the tetracyclines had magnesium ions coordinated in the SBS of *E. coli* NPET. These structural insights demonstrate that the mismatch bp, *U1782:U2586* in *E. coli* and C1965:C2768 in *C. acnes*, is the key determinant of the binding pose of the tetracyclines at the SBS.

The SBS of tetracyclines in the *C. acnes* and *E. coli* NPET reported here overlaps with that of the polyketide Tetracenomycin X. Unlike tetracyclines, Tetracenomycin X does not bind the 30S ribosomal subunit and exerts its translation inhibition exclusively via its NPET binding site. Recent structural studies on Tetracenomycin X demonstrated the proximity of the Tetracenomycin X binding site to the PTC causes conformational changes in the P-site aminoacyl-tRNA, inhibiting peptide bond formation *(12, 13)*. Overlaying the structures of Tetracenomycin X with the structures presented here shows that all three tetracyclines would likely cause similar conformational changes in the P-site aminoacyl-tRNA and inhibit the PTC (Fig. S3B). Interestingly, the efficacy of Tetracenomycin X was severely lowered by mutating the *U1782:U2586* of *E. coli* 70S ribosome to a C:C mismatch base pair as observed in *C. acnes* and *Thermus thermophilus* 70S ribosome*(12)*. In our structures of *C. acnes* and *E. coli* 70S ribosomes, containing a C:C and a U:U mismatch bp at the SBS, respectively, all three tested tetracyclines were able to bind both these bacterial ribosomes, albeit with characteristic differences in their binding poses (Fig. 1C-D and S2).

### Doxycycline dimerizes in the NPET of bacterial 70S ribosome

Doxycycline is unique in its ability to dimerize and pack the NPET of *C. acnes* 70S ribosome. We observed two dimers of Doxycycline bound stably in the NPET (Fig. 2A and S2A), in addition to the monomeric NPET SBS shared with Minocycline and Sarecycline (Fig. 1C-D). The first Doxycycline dimer (Dox-D1) stacks directly on the monomeric Doxycycline bound in the NPET SBS, creating a trimer of Doxycycline molecules close to the PTC. The second Doxycycline dimer (Dox-D2) binds near the solvent side of the NPET, stabilized by the 23S rRNA, uL22, and uL23. Both Doxycycline dimers are symmetrical and involve two magnesium ions (Fig. 2B). In the Doxycycline dimer, ring A is puckered at a 60° angle relative to rings B, C, and D, and the two monomers stack against each other with ring C of both monomers overlaying on top of each other. The planar axes of the two monomers are parallel to each other but offset by almost 90°, leading to the C6-methyl modification of ring C from one monomer stacking on to ring D of the other monomer (Fig. 2B). These stacking interactions are further stabilized by two magnesium ions that have a symmetrical coordination geometry within the Doxycycline dimer. Each magnesium ion is coordinated by the C11-carbonyl (ring C) and C12-hydroxyl (ring B) of one monomer with the C1-carbonyl and C2-carboxyamide group of ring A from the second monomer (Fig. 2B).

**Figure 2.**
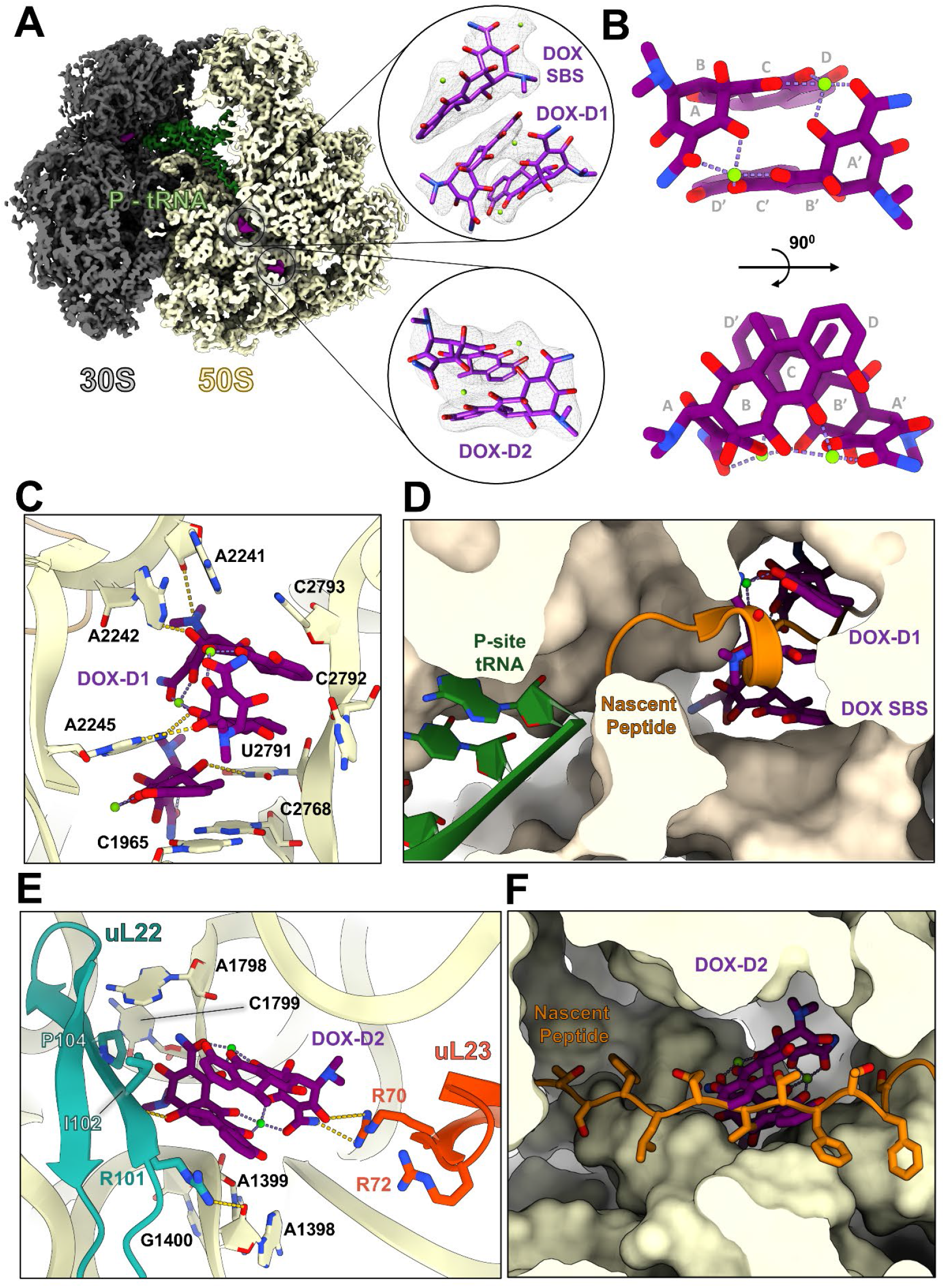
Doxycycline dimerizes in the NPET of bacterial 70S ribosome. **(A)** Cryo-EM reconstruction of *C. acnes* 70S ribosome bound to Doxycycline is shown. The insets highlight the atomic model and the corresponding cryo-EM density of Doxycycline SBS and the two Doxycycline dimers, Dox-D1 and Dox-D2, bound in the NPET. **(B)** A magnified view of the symmetrical Doxycycline dimer highlighting the critical interaction motifs. **(C)** Detailed view of the SBS of Doxycycline and Dox-D1 in the CA70S ribosome. **(D)** The structure of Doxycycline bound EC70S was aligned with that of EC70S with a nascent peptide chain (PDB:7ZP8) to highlight the steric clash of Doxycycline SBS and Dox-D1 with the nascent peptide. **(E)** Detailed view of the Dox-D2 binding site in the CA70S ribosome. **(F)** The structure of Doxycycline bound EC70S was aligned with that of EC70S with a nascent peptide chain (PDB:7ZP8) to highlight the steric clash of Dox-D2 with the nascent peptide.

Dox-D1 stacks on top of the monomeric Doxycycline bound in the SBS, creating a triple stack of Doxycycline molecules (Fig. 2C, S2A, and S2C). The Doxycycline dimer stacks against the 23S rRNA nucleotides A2241 *(A2058)*, A2242 *(A2059)*, and C2793 *(C2611)* that form the roof of the NPET, and A2245 *(A2062)* and C2792 *(C2610)* that form the sides of the NPET near the PTC (Fig. 2C). Most of these 23S rRNA nucleotides are part of the highly conserved PTC ring and are critical for the catalysis of peptide bond formation *(20)*. Therefore, interaction of Dox-D1 with these critical residues would most likely inhibit the PTC. Aligning the structure of a translating *E. coli* 70S ribosome with a nascent peptide (PDB:7ZP8) *(21)* to our structure of *E. coli* 70S ribosome bound to Doxycycline revealed that the three Doxycycline molecules completely occlude the NPET near the PTC and sterically clash with the nascent polypeptide chain (Fig. 2D).

Dox-D2 binds near the solvent side of the NPET, stacking between A1399 *(C1323)*, A1798 *(A1614)*, and C1799 *(C1615)* of the 23S rRNA, and further stabilized in the NPET by uL22 and uL23 (Fig. 2E). uL22 and uL23 are critical for ribosome biogenesis and play integral roles in protein translation. The β-hairpin loop of uL22 (G96-S117 / *G79-T100)* is the largest contributor of ribosomal protein interface for the NPET in bacteria and has been shown to be essential for proper folding of the nascent chain *(22)*. Dox-D2 binds at the interface of this uL22 β-hairpin loop and the 23S rRNA nucleotides A1398-G1400 *(A1322-G1324)* and A1798-A1800 *(A1614-A1616)*.

Most of the interactions for Dox-D2 binding come from one molecule of the dimer. Rings B and C of this Doxycycline molecule stack against A1398-G1400 *(A1322-G1324)* whereas ring A stacks against A1798-A1800 *(A1614-A1616)*. C6-methyl from this Doxycycline molecule of Dox-D2 stacks against R101 *(R84)* and I102 *(I85)* of uL22, while the neighboring C5-hydroxyl forms a hydrogen-bond with the peptide backbone between R101 *(R84)* and I102 *(I85)* (Fig. 2E). The second Doxycycline molecule of Dox-D2 provides additional interactions through a hydrogen bonding network between its C2-hydroxyl and C3-carboxyamide group and R70 *(R69)* of the β-hairpin uL23 loop (K67-P79 / *K66-S78)*. This loop of uL23 has been shown to be essential for sensing by the signal recognition particle that delivers a quarter of all proteins to the bacterial membrane for co-translational insertion *(23)*. Thus, Dox-D2 interactions with the β-hairpin loops of uL22 and uL23 in the NPET would impair their canonical functions in bacterial protein translation. Similar to Dox-D1 binding near the PTC, Dox-D2 also leads to steric clashes with the nascent peptide (Fig. 2F). We also observed a seventh molecule of Doxycycline bound to the 23S rRNA of the *C. acnes* 70S ribosome, intercalated between U1349 and G1350 (Fig. S4A). Minocycline was also observed at this binding site in the *C. acnes* 70S ribosome (Fig. S4B), but no binding was detected for Sarecycline (Fig. S4C), most probably due to steric clashes caused by its bulky C7 modification. This binding site is absent in the *E. coli* 70S ribosome due to sequence differences that alter the fold of the 23S rRNA in this region (Fig. S4D-E). Since this binding site is not close to either active center of the *C. acnes* ribosome, its impact on protein translation is likely minor. Overall, our structures predict that the multi-pronged translation inhibition by Doxycycline would be more potent than Minocycline and Sarecycline.

### Occupancy analysis reveals the concentration dependence of the tetracycline SBS

Depending on the mode of administration and the tissue type, the local concentration of tetracycline can vary significantly. To understand the effect of concentration on protein translation inhibition, we tested bacterial translation in the presence of increasing concentrations of Sarecycline, Minocycline, and Doxycycline. As expected, protein translation was inhibited more potently as the concentration of the tetracyclines was increased. Surprisingly, at higher concentrations (500 µM), Doxycycline was 2-3-fold more potent than Sarecycline and Minocycline (Fig. 3A). At lower concentrations (<200 µM), this difference was less pronounced. To understand this concentration dependency, we resolved cryo-EM reconstructions of *E. coli* 70S ribosome bound to Sarecycline, Minocycline, and Doxycycline at varying concentrations (80, 200, and 500 µM). To quantitatively measure the occupancy of tetracyclines in the CBS and SBS, we utilized resampled charge density (CD) maps (see Methods section for details) and calculated CD ratios between the tetracycline and neighboring RNA backbone atoms. This allowed us to calculate the relative occupancy of the tetracyclines at the CBS and SBS. The occupancy analysis revealed that the SBS of tetracyclines is a lower affinity binding site compared to the CBS and at lower concentrations, the tetracyclines bound only in the CBS. At a concentration of 200 µM and above, binding at both sites was observed. Similarly, the two Doxycycline dimer binding sites, Dox-D1 and Dox-D2, were occupied only at concentrations of 200 µM and above (Fig. S5A). Among the three tetracyclines tested here, Doxycycline showed the highest occupancy at the SBS (Fig. 3B and Table S3). Since the Dox-D1 dimer stacks on to the monomeric Doxycycline molecule in the SBS, it is likely that the additional stability provided by the stacking interactions with Dox-D1 leads to the higher occupancy of Doxycycline in the SBS. The presence of Doxycycline dimers in the NPET and the higher occupancy in the SBS at concentrations of 200 µM and above correlates well with the protein translation inhibition curve of Doxycycline (Fig. 3A). Thus, at lower concentrations, all three tetracyclines are equally effective due to binding only at the CBS. However, at higher concentrations, Doxycycline has enhanced potency in inhibiting translation due to its higher occupancy at the SBS and blocking the NPET by dimerization at two different sites.

**Figure 3.**
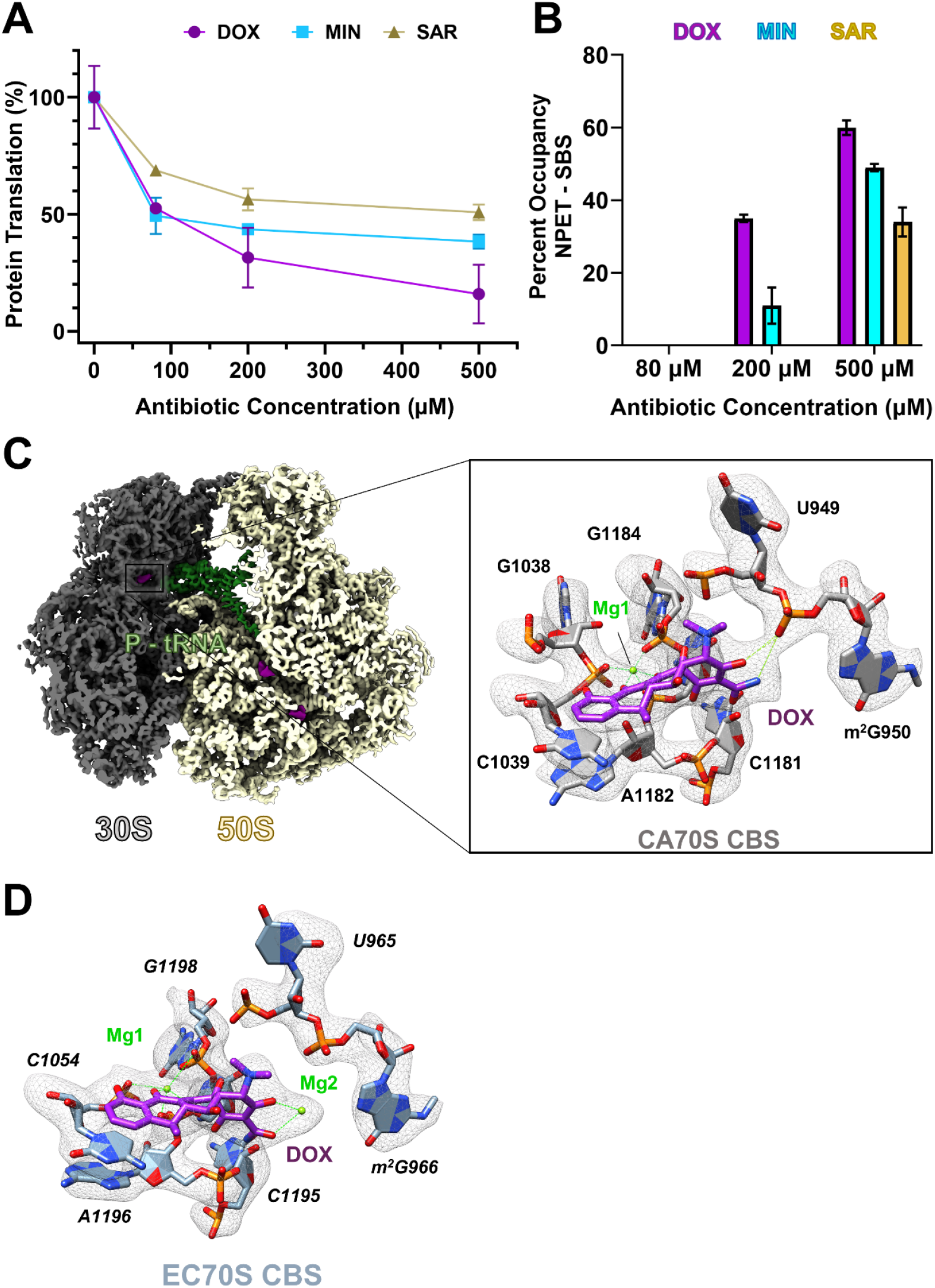
Differences in SBS Occupancy and CBS of Tetracyclines. **(A)** Inhibition of E. coli protein translation was tested for increasing concentrations of DOX, MIN, and SAR. The percent inhibitory effect is plotted against the corresponding concentration of the respective tetracycline. Relative occupancy of DOX, MIN, and SAR in the EC70S SBS at 80µM, 200µM, and 500µM is plotted in bar chart form. **(C-D)** A magnified view of the Doxycycline canonical binding site (CBS) for *C. acnes* (C) and *E. coli* (D) 70S ribosome is shown. The structural model is shown in stick format along with the corresponding cryo-EM map density in mesh. The tetracycline CBS for *E. coli* 70S ribosome involves two magnesium ions whereas for the *C. acnes* 70S ribosome, the second magnesium (Mg2) is absent and the C2-carboxyamide group of tetracycline forms a direct interaction with the phosphate backbone of m^2^G950. This trend was observed for all three tetracyclines tested.

### Differences in the CBS of Tetracyclines

The binding of all three tetracyclines at the CBS of *C. acnes* and *E. coli* 70S ribosome was consistent with previously reported structural studies *(6-9, 11)*. All three tetracyclines bind the 30S ribosomal subunit between h34 and h31 of the 30S head, above the mRNA channel and overlap with the binding site of the A-site tRNA anticodon loop (Fig. 3C-D). The CBS for tetracyclines involves an extensive hydrogen-bonding network with the phosphate backbone of the 16S rRNA. The sole stacking interaction in the CBS is provided by C1039 *(C1054)* of h34 that forms a three-ring pi stack with ring D of the tetracycline and A1182 *(A1196)*. As previously reported, a magnesium ion is coordinated by the hydroxyl groups of ring B and C of tetracycline and the phosphate backbone of C1039 *(C1054)*, A1183 *(A1197)*, and G1184 *(G1198)*. A notable difference between the CBS of tetracyclines for *C. acnes* and *E. coli* 70S ribosome is the coordination of a second magnesium ion. In the CBS of *E. coli* 70S ribosome, a second magnesium ion is coordinated by the C2-carboxyamide group and C3-hydroxyl group of tetracycline A ring along with the phosphate backbone of *2-methyl G966*. For *C. acnes* 70S ribosome, this second magnesium ion is missing and instead C3-hydroxyl of the tetracyclines is within hydrogen-bonding distance of the phosphate backbone of 2-methyl G950 *(2-methyl G966)*. This difference is observed for all the three tetracyclines tested, emphasizing species-specific ribosomal variation.

## Discussion

Our results establish that tetracyclines target both active centers of the bacterial ribosome, the mRNA decoding center and the PTC/NPET, to inhibit bacterial protein translation. This raises the question of why the SBS of tetracyclines was not observed in previous structural studies of tetracycline. A possible reason for this discrepancy is the method for structural characterization; almost all prior studies for characterizing tetracycline binding to bacterial ribosomes relied on X-ray crystallography and usually involved “soaking in” of the antibiotic into preformed crystals of the bacterial ribosome. Our comparative structural characterization of Sarecycline with *T. thermophilus* 70S ribosomes with X-ray crystallography and cryo-EM showed that the SBS for Sarecycline was only observed via cryo-EM (Fig. S6). This may be due to solubility issues of the drug under crystallization conditions or potential solvent accessibility limitations for the ribosomal NPET in a preformed crystal lattice.

A major determinant of the tetracycline SBS is the non-canonical Watson-Crick base pair *U1782:U2586*, and further stabilization by *A2062*. The structural framework of the tetracycline SBS is conserved across bacterial species, with the universal presence of the *A2062* residue and the U:U or C:C mismatch base pair. Although we did not observe a correlation between the occupancy levels of tetracyclines at the SBS and the identity of the mismatch base pair (U:U vs C:C), we observed a correlation between the binding pose of the tetracyclines in the SBS (Fig. 1C-D and S2A-D). The binding pose of the tetracyclines at the SBS showed a ∼90° rotation for the U:U vs C:C non-canonical Watson-Crick base pair in the SBS (Figure S3A). This is most likely due to the difference in electrostatic potential created by a C:C base pair compared to a U:U base pair. Interestingly, this 90° rotation seems to cause a steric clash for the unique C7-extension of Sarecycline with *U2609* in the *E. coli* 70S ribosome and the drug is accommodated by flipping the drug by 180°. The occupancy of Sarecycline at the SBS is two-fold lower compared to Doxycycline, which lacks any modification at the C7 position, indicating that such modifications can be used to tailor future tetracycline derivatives to differentially target the CBS and SBS as well as exploit species-specific ribosomal differences.

Apart from bacterial ribosomes, a C:C mismatch base pair is also found in the exact same location in the human 55S mitochondrial ribosome, and recent studies demonstrated that Tigecycline, a third-generation tetracycline, can bind at this site and potently inhibit mito-ribosomal protein translation*(24, 25)*. Overlaying our tetracycline-bound structures of *C. acnes* 70S ribosome with the Tigecycline-bound human 55S mitochondrial ribosome revealed remarkable conservation in the binding site as well as the interaction network of the drug (Fig. 4A). Interestingly, the C9 extension of Tigecycline extends towards the PTC, creating steric clashes with the P-site aminoacyl-tRNA. Thus, adding bulkier modifications at the C8 and C9 positions of tetracycline that would project into the PTC presents an attractive strategy for designing the next generation of tetracyclines that can more efficiently target the bacterial PTC in addition to the mRNA decoding center.

**Figure 4.**
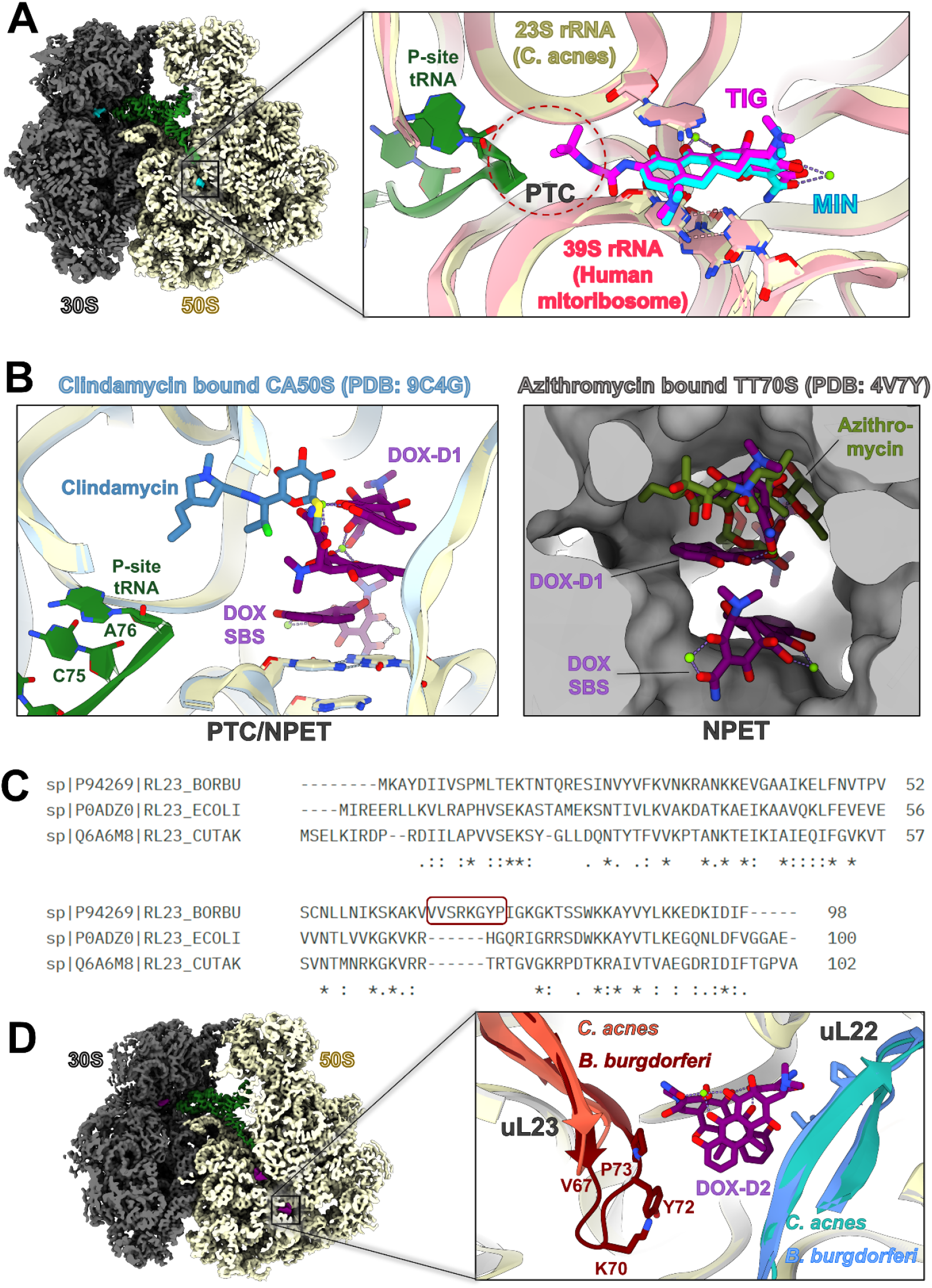
Structural insights for the development of future tetracyclines. **(A)** The structure of Minocycline bound CA70S was aligned with the structure of Tigecycline bound to 55S human mitochondrial ribosome (PDB: 8K2A). The inset shows a detailed view of the PTC/NPET, highlighting the conservation in the SBS binding site of tetracyclines across species. **(B)** The structure of Doxycycline bound CA70S was aligned with the structure of Clindamycin bound to 50S subunit of *C. acnes* (PDB:9C4G) (left panel), and the structure of Azithromycin bound *Thermus thermophilus* (TT) 70S ribosome (PDB:4V7Y) (right panel). Both panels highlight the overlap of the binding site for Dox-D1 with that of lincosamides and macrolides, respectively. **(C)** Sequence alignment of uL23 protein from *Borrelia burgdorferi* (Uniprot: P94269), *C. acnes* (Uniprot: Q6A6M8), *and E. coli* (Uniprot: P0ADZ0) was carried out using Clustal Omega. The unique insertion observed in *Borrelia burgdorferi* uL23 is highlighted with a maroon box. **(D)** The structure of Doxycycline bound CA70S was aligned with the structure of hibernating 70S ribosome from *Borrelia burgdorferi* (PDB:8FMW). The inset shows the Dox-D2 binding site and potential enhanced interaction with the unique insertion of *Borrelia burgdorferi* uL23.

Doxycycline is one of the top five most-prescribed antibiotics in the United States, with over 20 million prescriptions annually *(26)*. Among the wide range of diseases treated by Doxycycline, it has emerged as the first line of treatment against tick-borne illnesses like Lyme disease and Rocky Mountain Spotted Fever. Our results show that among the three tetracyclines tested herein, Doxycycline was unique in its ability to form dimers and extensively block the NPET at multiple locations. The binding site of the first dimer, Dox-D1, overlaps with that of macrolides *(27)* and lincosamides *(9, 28)* (Fig. 4B). Dox-D1 stacks on top of the Doxycycline molecule bound at the SBS in the NPET, and this trimer of Doxycycline molecules completely blocks the NPET near the PTC. The second dimer, Dox-D2, binds in the NPET, further away from the PTC, and is a unique binding site among antibiotics targeting the bacterial ribosome. Comparing the structure of the 70S ribosome from the Lyme disease-causing bacteria *Borrelia burgdorferi (Bbu)* with our Doxycycline bound 70S ribosomal structures revealed a more favorable Dox-D2 binding site in *Bbu* 70S ribosome (Fig. 4C) compared to *C. acnes* or *E. coli* 70S ribosome. This difference arises from the unique extension in the uL23 loop of *Bbu* 70S ribosome that extends into the NPET. The extended uL23 loop in *Bbu* 70S ribosome places Y72 and P73 of uL23 as additional binding interactions for Dox-D2 in addition to the conserved interaction network provided by uL22 and 23S rRNA (Fig. 4D). This may provide a structural rationale for Doxycycline’s efficacy and clinical utility in treating Lyme disease, although direct comparative studies with other tetracyclines are limited.

The rising wave of antimicrobial resistance is one of the most daunting challenges facing public health and necessitates the urgent need for development of novel antibiotics. The tetracycline class of antibiotics is widely used to treat an array of bacterial infections and newer generation of tetracyclines are being developed to enhance their efficacy and overcome bacterial resistance mechanisms. The in-depth structural characterization of tetracyclines carried out herein redefines the mechanism of action for tetracyclines, wherein both the mRNA decoding center (CBS) and the PTC/NPET (SBS) of the bacterial ribosome are targeted. Remarkably, the interaction network was well-conserved for all three tested tetracyclines at both the CBS and SBS, and this dual-targeting mechanism was observed across bacterial species. Our findings provide a structural framework for the rational design of the next-generation of narrow-spectrum tetracyclines that can leverage species-specific differences observed in the 70S ribosomes from pathogenic bacteria and tackle the growing threat of antimicrobial resistance.

## Author contributions

Conceptualization: SCD, IBL, CGB

Methodology: SCD, IBL, JW

Investigation: SCD, IBL, JW

Funding acquisition: CGB

Writing – original draft: SCD

Writing – review & editing: SCD, IBL, JW, and CGB

## Data Availability

The coordinates and cryo-EM density maps for all the structures reported here have been deposited in the PDB and EMDB, respectively. The accession codes are as follows – *E. coli* 70S ribosome bound to Doxycycline (PDB: 9PIH, EMD-71667); *E. coli* 70S ribosome bound to Sarecycline (PDB: 9PII, EMD-71668); *E. coli* 70S ribosome bound to Minocycline (PDB: 9PIJ, EMD-71669); *C. acnes* 70S ribosome bound to Doxycycline (PDB: 9PJ7, EMD-71682); *C. acnes* 70S ribosome bound to Sarecycline (PDB: 9PJ8, EMD-71683); *C. acnes* 70S ribosome bound to Minocycline (PDB: 9PJ9, EMD-71684).

## Conflict of Interest

CGB has served as an investigator and/or consultant for Almirall, Ortho Dermatologics, Sun Pharma, and Teladoc.

### Acknowledgements and Funding

We acknowledge the Yale cryo-EM core facility, and the Laboratory for BioMolecular Structure (LBMS) at Brookhaven National Laboratory, supported by the DOE Office of Biological and Environmental Research (KP1607011), for access to their Krios cryo-EM. This work was funded by research grants from Almirall, Ortho Dermatologics, and the American Acne and Rosacea Society (to CGB).

## List of Supplementary Materials

1. Materials and Methods
2. Supplementary Figure S1-S6
3. Supplementary Table S1-S3

## References

1. B. M. Duggar, Aureomycin; a product of the continuing search for new antibiotics. Ann N Y Acad Sci 51, 177–181 (1948).

2. H. E. Baldwin, D. B. Ward, Jr., Fifty Years of Minocycline and Its Evolution: A Dermatological Perspective. J Drugs Dermatol 20, 1031–1036 (2021).

3. B. A. Cunha, C. M. Sibley, A. M. Ristuccia, Doxycycline. Ther Drug Monit 4, 115–135 (1982).

4. E. D. Deeks, Sarecycline: First Global Approval. Drugs 79, 325–329 (2019).

5. Y. R. Lee, C. E. Burton, Eravacycline, a newly approved fluorocycline. Eur J Clin Microbiol Infect Dis 38, 1787–1794 (2019).

6. D. E. Brodersen et al., The structural basis for the action of the antibiotics tetracycline, pactamycin, and hygromycin B on the 30S ribosomal subunit. Cell 103, 1143–1154 (2000).

7. L. Jenner et al., Structural basis for potent inhibitory activity of the antibiotic tigecycline during protein synthesis. Proc Natl Acad Sci U S A 110, 3812–3816 (2013).

8. A. I. Cocozaki et al., Resistance mutations generate divergent antibiotic susceptibility profiles against translation inhibitors. Proc Natl Acad Sci U S A 113, 8188–8193 (2016).

9. H. Paternoga et al., Structural conservation of antibiotic interaction with ribosomes. Nat Struct Mol Biol 30, 1380–1392 (2023).

10. Z. Zhang, C. E. Morgan, R. A. Bonomo, E. W. Yu, Cryo-EM Determination of Eravacycline-Bound Structures of the Ribosome and the Multidrug Efflux Pump AdeJ of Acinetobacter baumannii. mBio 12, e0103121 (2021).

11. I. B. Lomakin, S. C. Devarkar, S. Patel, A. Grada, C. G. Bunick, Sarecycline inhibits protein translation in Cutibacterium acnes 70S ribosome using a two-site mechanism. Nucleic Acids Res 51, 2915–2930 (2023).

12. I. A. Osterman et al., Tetracenomycin X inhibits translation by binding within the ribosomal exit tunnel. Nat Chem Biol 16, 1071–1077 (2020).

13. E. C. Leroy, T. N. Perry, T. T. Renault, C. A. Innis, Tetracenomycin X sequesters peptidyl-tRNA during translation of QK motifs. Nat Chem Biol 19, 1091–1096 (2023).

14. K. N. Agwuh, A. MacGowan, Pharmacokinetics and pharmacodynamics of the tetracyclines including glycylcyclines. J Antimicrob Chemother 58, 256–265 (2006).

15. B. A. Cunha, Oral doxycycline for non-systemic urinary tract infections (UTIs) due to P. aeruginosa and other Gram negative uropathogens. Eur J Clin Microbiol Infect Dis 31, 2865–2868 (2012).

16. G. Campistron et al., Pharmacokinetics and bioavailability of doxycycline in humans. Arzneimittelforschung 36, 1705–1707 (1986).

17. A. S. Malmborg, Bioavailability of doxycycline monohydrate. A comparison with equivalent doses of doxycycline hydrochloride. Chemotherapy 30, 76–80 (1984).

18. J. Paik, Topical Minocycline Foam 4%: A Review in Acne Vulgaris. Am J Clin Dermatol 21, 449–456 (2020).

19. Z. Batool, I. B. Lomakin, Y. S. Polikanov, C. G. Bunick, Sarecycline interferes with tRNA accommodation and tethers mRNA to the 70S ribosome. Proc Natl Acad Sci U S A 117, 20530–20537 (2020).

20. A.E. d’Aquino et al., Mutational characterization and mapping of the 70S ribosome active site. Nucleic Acids Res 48, 2777–2789 (2020).

21. M. Ahn et al., Modulating co-translational protein folding by rational design and ribosome engineering. Nat Commun 13, 4243 (2022).

22. M. G. Lawrence et al., The extended loops of ribosomal proteins uL4 and uL22 of Escherichia coli contribute to ribosome assembly and protein translation. Nucleic Acids Res 44, 5798–5810 (2016).

23. K. Denks et al., The signal recognition particle contacts uL23 and scans substrate translation inside the ribosomal tunnel. Nat Microbiol 2, 16265 (2017).

24. X. Li et al., Structural basis for differential inhibition of eukaryotic ribosomes by tigecycline. Nat Commun 15, 5481 (2024).

25. Q. Shao et al., T cell toxicity induced by tigecycline binding to the mitochondrial ribosome. Nat Commun 16, 4080 (2025).

26. Centers for Disease Control and Prevention, National Center for Emerging and Zoonotic Infectious Diseases (NCEZID), Division of Healthcare Quality Promotion (DHQP), Outpatient Antibiotic Prescriptions — United States, 2022 (2022 https://archive.cdc.gov/#/details? url= https://www.cdc.gov/antibiotic-use/data/report-2022.html).

27. D. Bulkley, C. A. Innis, G. Blaha, T. A. Steitz, Revisiting the structures of several antibiotics bound to the bacterial ribosome. Proc Natl Acad Sci U S A 107, 17158–17163 (2010).

28. I. B. Lomakin, S. C. Devarkar, A. Grada, C. G. Bunick, Mechanistic Basis for the Translation Inhibition of Cutibacterium acnes by Clindamycin. J Invest Dermatol 144, 2553–2561 e2553 (2024).

