## Supplementary Figures, Tables, and Methods for "Structures redefine the mechanism of action for tetracyclines"

### Materials and Methods

#### Antibiotics Used for Biochemical and Structural Studies

Doxycycline and Minocycline were purchased from Millipore Sigma, USA (Cat. # D9891 and M9511, respectively). Sarecycline was kindly provided by Almirall, LLC.

#### Cell growth and ribosome purification.

*Cutibacterium acnes* cells were grown in anaerobic conditions using nitrogen and $CO_2$ generators (BD GasPack EZ with indicator, cat. # 260001). *C. acnes* cells (ATCC® 11827™) were reactivated and plated according to ATCC instruction on blood agar contact plates (REMEL™, cat. # R111007). Cells from one plate were transferred to 6 liters flask filled with two liters of Brain Heart Infusion Broth (OXOID, cat. # CM1135) and were grown for 40 hours at 37°C in shaker at 100 rpm. Identity of cells was confirmed by 16S Sanger Sequencing (CD Genomics, USA).

Ribosomes were purified from 5 g frozen cells. Cells were suspended in 50 ml of buffer B: 20 mM HEPES-KOH, pH 7.5, 200 mM KCl, 20 mM MgAc, 1 mM DTT, 1 mg/mL lysozyme, 1 tablet of cOmplete™ Protease Inhibitor Cocktail (Roche) and 100 U of DNaseI (RNase free). Cells were lysed using microfluidizer at 15000 psi for 3 cycles and then centrifugated at 18000 rpm, 4°C for 30 min in Type 45 Ti rotor (Beckman Coulter). Supernatant was layered onto 25 ml sucrose cushion (20 mM HEPES-KOH, pH 7.5, 500 mM KCl, 20 mM $MgCl_2$, 1 mM DTT, 1.1 M sucrose) and centrifuged at 42000 rpm, 4°C for 21 hours in the same rotor. Ribosomal pellets were suspended in high-salt buffer (20 mM HEPES-KOH, pH 7.5, 500 mM KCl, 10 mM MgAc, 1 mM DTT) by gentle shaking for 3 hours at 4°C and then layered again on the sucrose cushion as described. Ribosomal pellet was suspended in high-salt buffer and then layered onto sucrose gradient (20 mM HEPES-KOH [pH 7.5], 60 mM KCl, 10 mM $MgCl_2$, 1 mM DTT, 10%–40% [w/v] sucrose) to resolve 70S ribosomes by ultracentrifugation at 20000 rpm, 4°C for 16 hours in SW 32 Ti rotor (Beckman Coulter). Fractions containing 70S particles were concentrated in a spin concentrator (100,000 MWCO [molecular weight cutoff], Amicon®) and buffer-exchanged to a ribosome buffer (20 mM HEPES-KOH, pH 7.5, 60 mM KCl, 10 mM $MgCl_2$, 1 mM DTT) prior to flash-freezing in liquid nitrogen.

70S ribosomes from *E. coli* were derived from PURExpress® Δ Ribosome Kit (New England Biolabs, USA. Cat. # E3313S) and used according to the manufacturer's protocol.

#### Preparation of the reassociated 70S ribosomes.

After centrifugation through the second sucrose cushion (see above) the pellet was resuspended in dissociation buffer (20mM Tris HCl, pH 7.4, 60 mM $NH_4Cl$, 1 mM $MgCl_2$, 0.5 mM EDTA, 14 mM β-mercaptoethanol (βME)), layered onto 15-40% sucrose gradient to resolve 30S and 50S subunits (20 mM HEPES-KOH [pH 7.5], 60 mM $NH_4Cl$, 1 mM MgCl2, 14 mM βME, 15%–40% [w/v] sucrose) by centrifugation at 28000 rpm, 4°C for 16 hours in the SW 32 Ti rotor. Fractions corresponding to 50S and 30S subunits were combined, transferred to the reassociation buffer (20mM Tris HCl, pH 7.4, 60 mM NH4Cl, 10 mM MgCl2, 0.5 mM EDTA, 14 mM βME) by concentration-dilution (repeated twice) using a spin concentrator. Ribosomes were incubated at 37°C for one hour, and then were layered onto 10-40% sucrose gradient to resolve 70S subunit and collected as described above.

**Ribosomal complex formation**

Reaction was done in a total volume of 20 μl with the final concentration of 0.125 μM for the reassociated 70S *C. acnes* or *E. coli* ribosomes, 5.0 μM for mRNA (32MF), 500 μM for the specified tetracycline (unless stated otherwise), and a buffer composed of 20 mM Tris HCl, pH 7.4, 60 mM NH4Cl, 10 mM $MgCl_2$, 0.5 mM EDTA, 10 mM βME at 37°C for 30 minutes. Then $tRNA_i^{fMet}$ was added to a final concentration of 2.5 μM and the incubation was continued for another 30 minutes.

**Cryo-EM sample preparation, data collection and processing**

The assembled ribosomal complexes (4 μl) were applied on a 300 mesh C-Flat R2/1 holey carbon grids (Electron Microscopy Sciences, USA) pre-treated by glow-discharging at 11 mA for 30 s. The grids were blotted (blot force 1, blot time 7 s) at 25°C with 100% humidity and plunge-frozen in liquid ethane using FEI Vitrobot Mark IV (Thermo Fisher, USA). CHAPSO (Millipore Sigma) was used at a final concentration of 0.05% to improve particle distribution and vitreous ice homogeneity.

Images were acquired on 300 kV FEI Titan Krios electron microscope (Thermo Fisher) equipped with a post-GIF Gatan K3 direct detector in super-resolution mode, at a nominal calibrated magnification of 81,000x and a physical pixel size of 1.07 Å. The energy filter slit width was set to 15 eV. The datasets with *E. coli* 70S ribosomes and lower concentrations of SAR, DOX, and MIN (80 μM and 200 μM) were collected on a 200 kV Glacios electron microscope (Thermo Fisher) equipped with a Gatan K3 direct detector in super-resolution mode, at a nominal calibrated magnification of 45,000x and a physical pixel size of 0.868 Å. Automated data collection was set up using either SerialEM (*1*) or EPU (ThermoFisher Scientific) software suites. Data were collected with a dose of 15 electrons per pixel per second and with a defocus range of -0.8 μm to -1.8 μm, and a total dose of ~50 e/$Å^2$.

Data processing procedures were carried out using standard pipelines in cryoSPARC v4 (Supplementary Figure S1, Table S1) (*2*). cryoSPARC 'Blob picker' was used on a subset of 500 micrographs and the selected particles were used for 2D classification to generate ideal templates. These templates were used to pick particles from the entire dataset using cryoSPARC's template picker. After 2D Classification, 3D reconstructions were generated using cryoSPARC's Ab Initio and Heterogenous Refinement jobs. Particles belonging to 3D classes representing junk particles and unbound 50S subunit were discarded and those belonging to 70S ribosome were carried forward. Further 3D classification yielded a final particle stack of 70S ribosomes with bound P-site tRNA, SAR/DOX/MIN, and mRNA. These particles were then polished using cryoSPARC's Reference Based Motion Correction job and used for generating the final reconstruction using Homogenous Refinement.

**Model building**

Initial atomic model and restraints for the *E. coli* and *C. acnes* 70S ribosome reconstructions were generated using PDB ID 7K00 and 8CRX, respectively (*3, 4*). The final models of the *E. coli* and *C. acnes* 70S ribosome in complex with mRNA, tRNA and SAR/DOX/MIN were generated by multiple rounds of model building in COOT (*5*), followed by refinement in PHENIX (*6*).

Atomic model coordinates and electron density maps were deposited in RCSB PDB and EMDB databases, respectively. All figures showing atomic models and cryo-EM map densities were generated using either UCSF Chimera (*7*) or UCSF ChimeraX (*8*).

**Antibiotic Occupancy Analysis**

Coordinates of bound antibiotic at the CBS or SBS and its interacting nucleotides within 10 Å were extracted and translated into the center of a cubic box with edge length of 40 Å for calculation of a reference map at 1.5Å resolution using CCP4 package (*9*). This reference map was used for resampling the experimental maps (typically around 2.5-2.8 Å resolution) after the maps were also translated by the same transformation matrix using Chimera (*10*). The coordinates were refitted into the resampled experimental maps before plotting the experimental data. For all three tetracycline antibiotics, 18 circularly connected atoms around the four rings were extracted for plotting. For nucleic acid backbones, 16 consecutively bonded atoms of three nucleotides that are nearest to the antibiotic were selected for plotting: C1'-C2'- C3'-O3' of the 5' nucleotide, and P-O5'-C5'-C4'-C3'-O3' of the two following nucleotides.

The resampled maps were Fourier inverted into structure factors in ascii format using CCP4 package (*9*) and were used for Fourier summation along the plotting axes as a function of an added ΔB Gaussian smoothening function. A ΔB increment was 10 $Å^2$ within 100 $Å^2$ and 25 $Å^2$ for the remaining range up to 500 $Å^2$. The experimental maps analyzed were CD maps, which were generated through Laplacian operation, rescaled with negation scale factor of -1 before being resampled, also using Chimera (*10, 11*). The mean CD ratios between antibiotic and RNA backbone plots were calculated and rescaled to the reference of Minocycline CBS.

***In vitro* translation and luciferase inhibition assays.**

These assays were used to measure *in vitro* ribosome activity via production of firefly luciferase in the PURExpress® *in vitro E. coli* Protein Synthesis Kit (New England Biolabs, USA, cat. # E6800S). Cell-free extract was prepared according to the manufacturer's instruction and supplemented with the lucRNA2caa10 mRNA (5 ng), SuperaseIN™ RNase inhibitor (2U, Ambion, cat. # AM2694) and water to a final volume of 8.3 µl and incubated at 37°C for 5 minutes. Then 2 µl of the indicated antibiotic were added to reach a final concentration of 80 µM, 200 µM, or 500 µM. The reaction proceeded for 2.5 hours at 37°C in the shaker and then transferred in the 96 well white plate. The luminescent signal was detected in 1-2 minutes after addition of 50 µl of the Luciferase Assay Reagent (Promega, USA, cat. # E1500) using multimode plate reader TriStar LB941 (Berthold technologies LTD, UK) and analyzed using Prism 9 (Version 9.3.1) software (GraphPad Software, LLC). All experiments were done in triplicate. Translation efficiency was calculated as normalized end-point values of luminescence with measurement interval time of 0.2 second.

### Figure S1

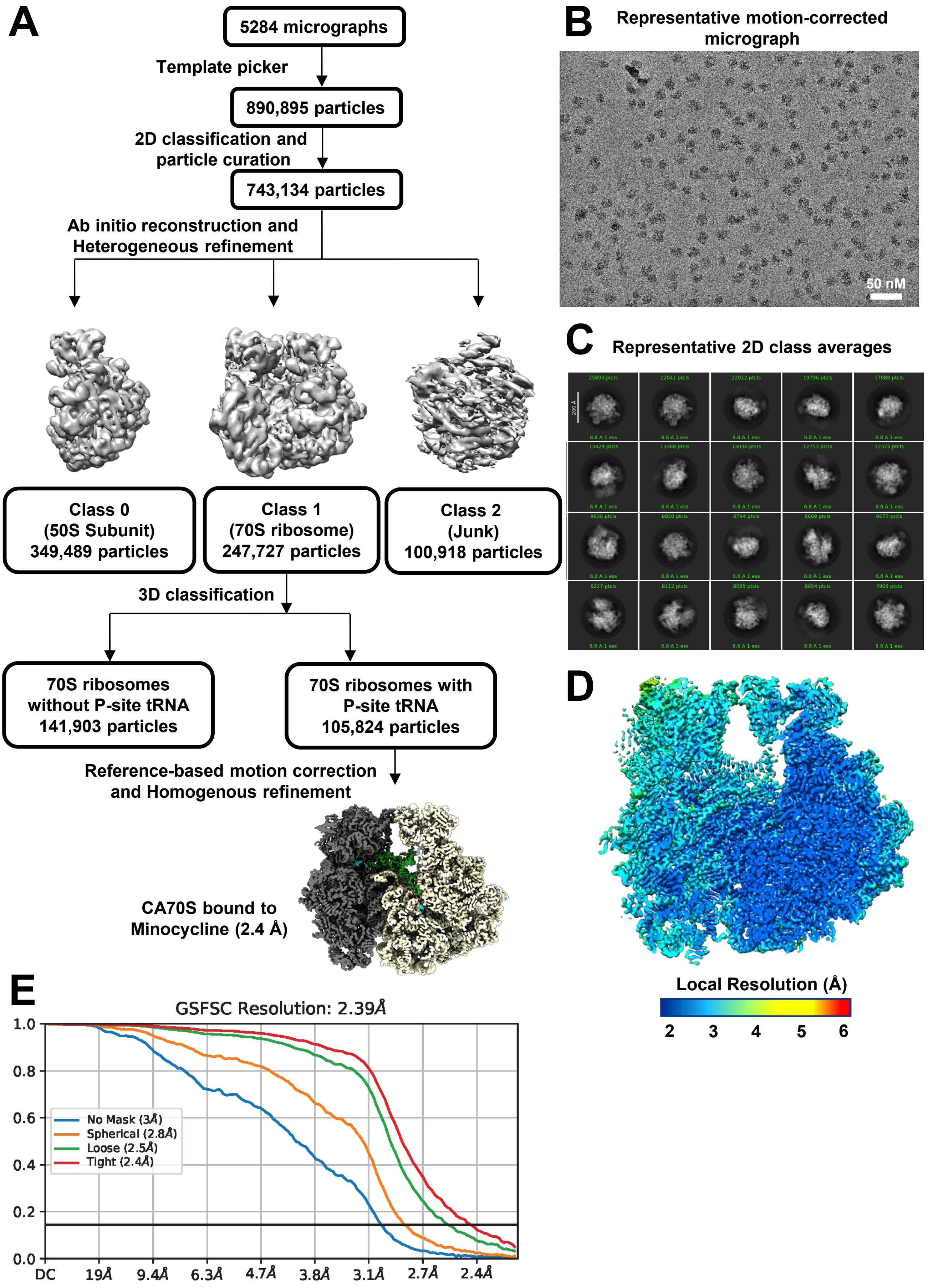
A
5284 micrographs
Template picker
890,895 particles
2D classification and particle curation
743,134 particles
Ab initio reconstruction and Heterogeneous refinement
Class 0 (50S Subunit) 349,489 particles
Class 1 (70S ribosome) 247,727 particles
Class 2 (Junk) 100,918 particles
3D classification
70S ribosomes without P-site tRNA 141,903 particles
70S ribosomes with P-site tRNA 105,824 particles
Reference-based motion correction and Homogenous refinement
CA70S bound to Minocycline (2.4 Å)
B
Representative motion-corrected micrograph
50 nM
C
Representative 2D class averages
D
Local Resolution (Å)
2
3
4
5
6
E
GSFSC Resolution: 2.39Å
No Mask (3Å)
Spherical (2.8Å)
Loose (2.5Å)
Tight (2.4Å)
1.0
0.8
0.6
0.4
0.2
0.0
DC
19Å
9.4Å
6.3Å
4.7Å
3.8Å
3.1Å
2.7Å
2.4Å

**Figure S1. (A)** The overall cryo-EM data processing workflow for the Minocycline bound *C. acnes* 70S (CA70S) ribosome reconstruction is shown. All other cryo-EM reconstructions reported as part of this study were obtained after following an identical workflow in cryoSPARC v4. **(B-C)** Representative motion-corrected micrograph **(B)** and 2D class averages **(C)** for the Minocycline-CA70S dataset are shown. **(D)** A cross-section view of the Minocycline-CA70S cryo-EM reconstruction highlighting the local resolution of the map in a color-coded format is shown. The local resolution varies from ~2 Å for the core regions to ~5 Å for the peripheral dynamic regions of rRNAs and ribosomal protein segments. **(E)** The gold-standard Fourier Shell Correlation (GS-FSC) curve for the Minocycline-CA70S reconstruction is shown.

#### Figure S2

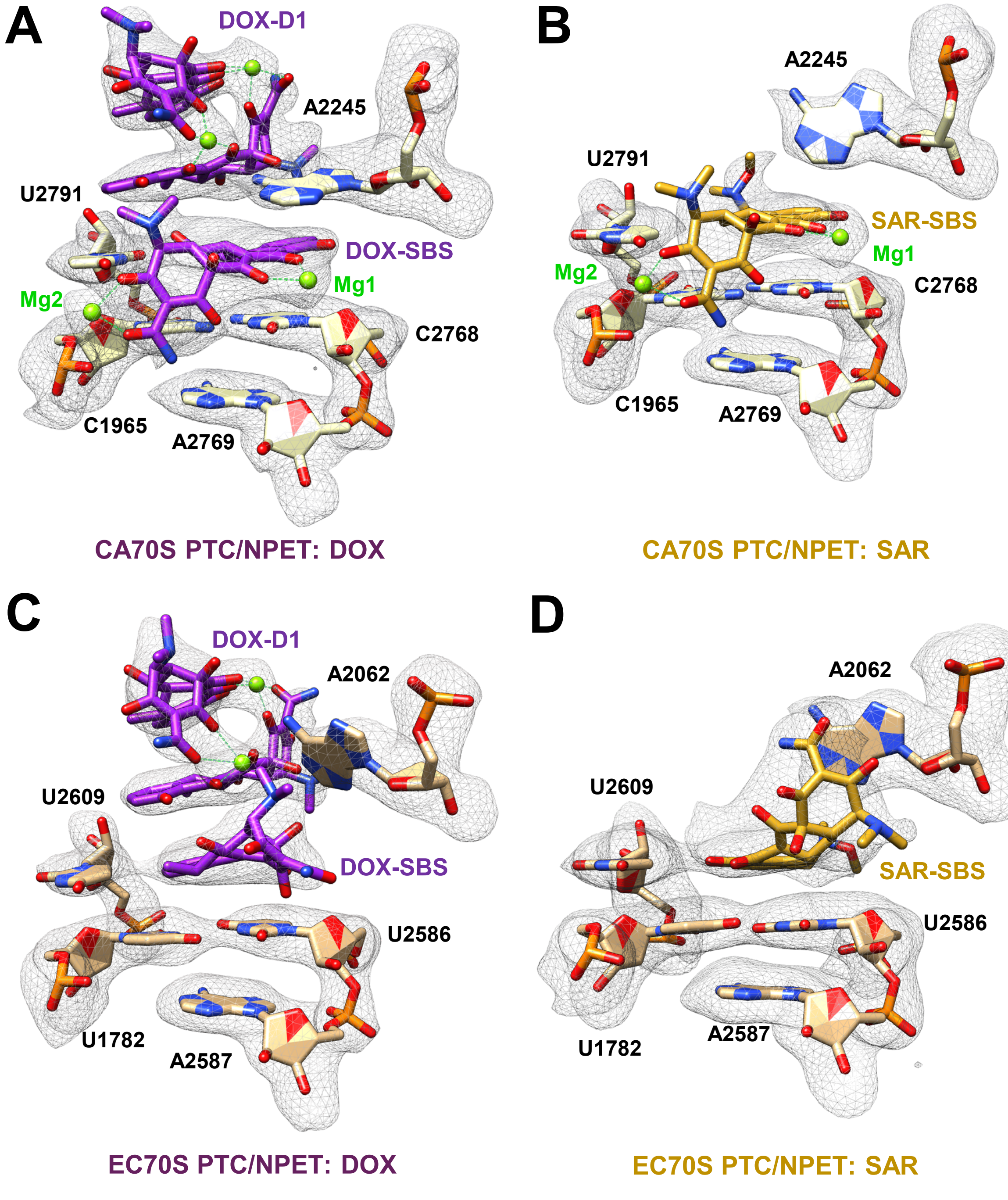


**Figure S2.** A magnified view of the Doxycycline **(A,C)** and Sarecycline **(B,D)** SBS for *C. acnes* 70S and *E. coli* 70S ribosome is shown. The structural model is shown in stick format along with the corresponding cryo-EM map density in mesh. Tetracyclines coordinate two magnesium ions (Mg1 and Mg2) in the SBS for *C. acnes* 70S ribosome but not for the *E. coli* 70S ribosome SBS. In contrast, the Doxycycline dimer (DOX-D1) involves two magnesium ions and has an identical binding pose in both *C. acnes* 70S and *E. coli* 70S ribosome.

### Figure S3

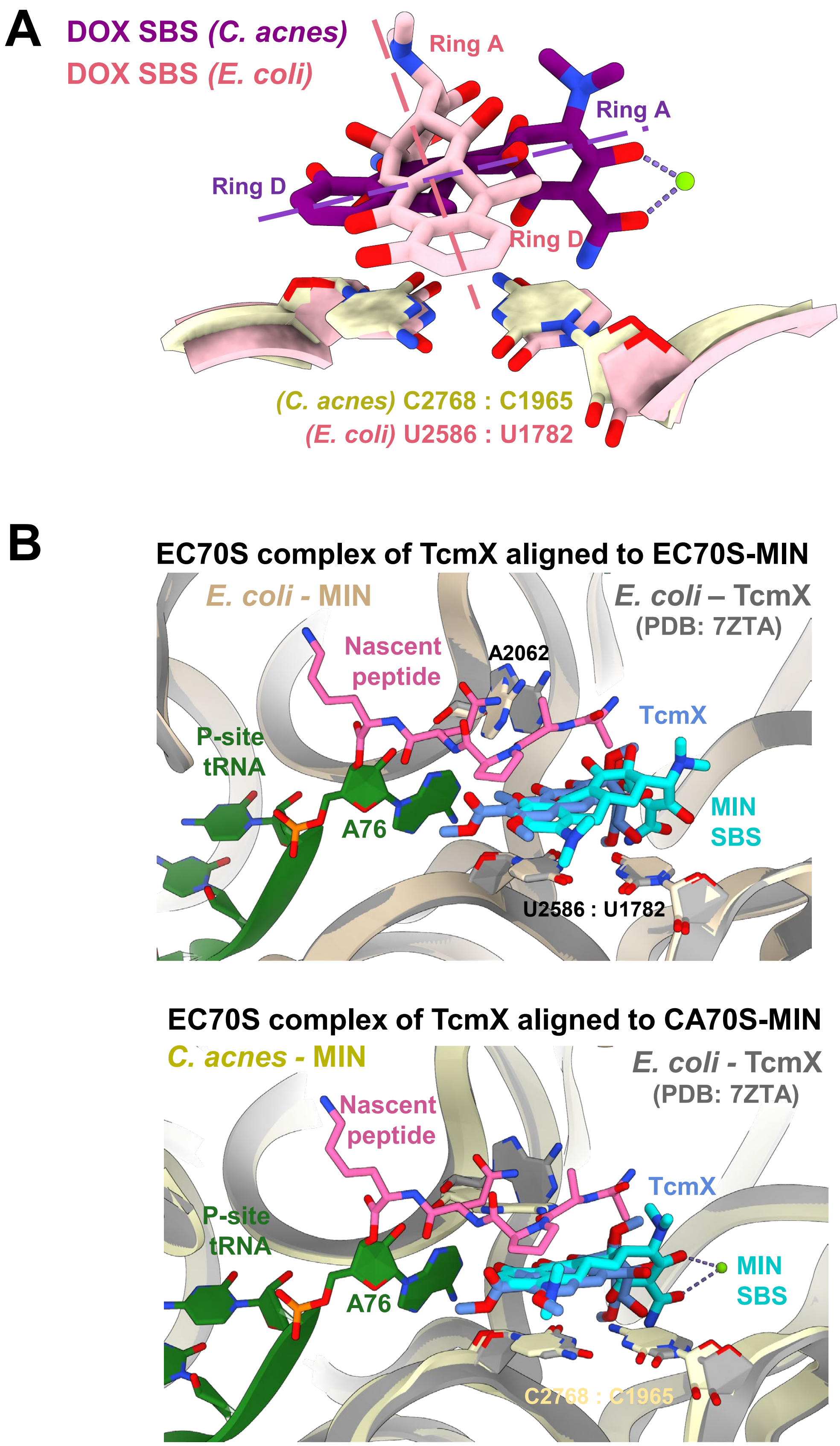
A
DOX SBS (C. acnes)
DOX SBS (E. coli)
Ring A
Ring A
Ring D
Ring D
(C. acnes) C2768 : C1965
(E. coli) U2586 : U1782
B
EC70S complex of TcmX aligned to EC70S-MIN
E. coli - MIN
E. coli – TcmX
(PDB: 7ZTA)
Nascent
peptide
A2062
TcmX
P-site
tRNA
MIN
SBS
A76
U2586 : U1782
EC70S complex of TcmX aligned to CA70S-MIN
C. acnes - MIN
E. coli - TcmX
(PDB: 7ZTA)
Nascent
peptide
TcmX
P-site
tRNA
MIN
SBS
A76
C2768 : C1965

**Figure S3. (A)** A magnified view of the Doxycycline SBS for *C. acnes* (purple) and *E. coli* (pink) 70S ribosome is shown. The non-canonical Watson-Crick base pairs C2768:C1965 (*C. acnes*) and U2586:U1782 (*E. coli*) are also shown to highlight the alignment of the 23S rRNA of *C. acnes* and *E. coli* 70S ribosome. The axis of Doxycycline in the SBS of *C. acnes* (purple) and *E. coli* (pink) 70S ribosome is shown with dotted lines to highlight the drastic change in the binding pose of Doxycycline induced by the difference in the non-canonical Watson-Crick base pair. A similar trend is observed for Sarecycline and Minocycline. **(B)** TcmX inhibits protein synthesis at QK motifs of the nascent peptide by stabilizing a non-productive conformation of the A76 nucleotide of peptidyl site aminoacyl-tRNA. The cryo-EM structure of TcmX bound to *E. coli* 70S ribosome paused at a QK motif of the nascent peptide (PDB: 7ZTA) was overlayed with Minocycline bound *E. coli* (top panel) and *C. acnes* (bottom panel) 70S ribosome structures reported herein. The binding pose of Minocycline and TcmX as well as the critical surrounding nucleotides belonging to the PTC ring overlay closely between these structures, suggesting a similar mechanism of action.

### Figure S4

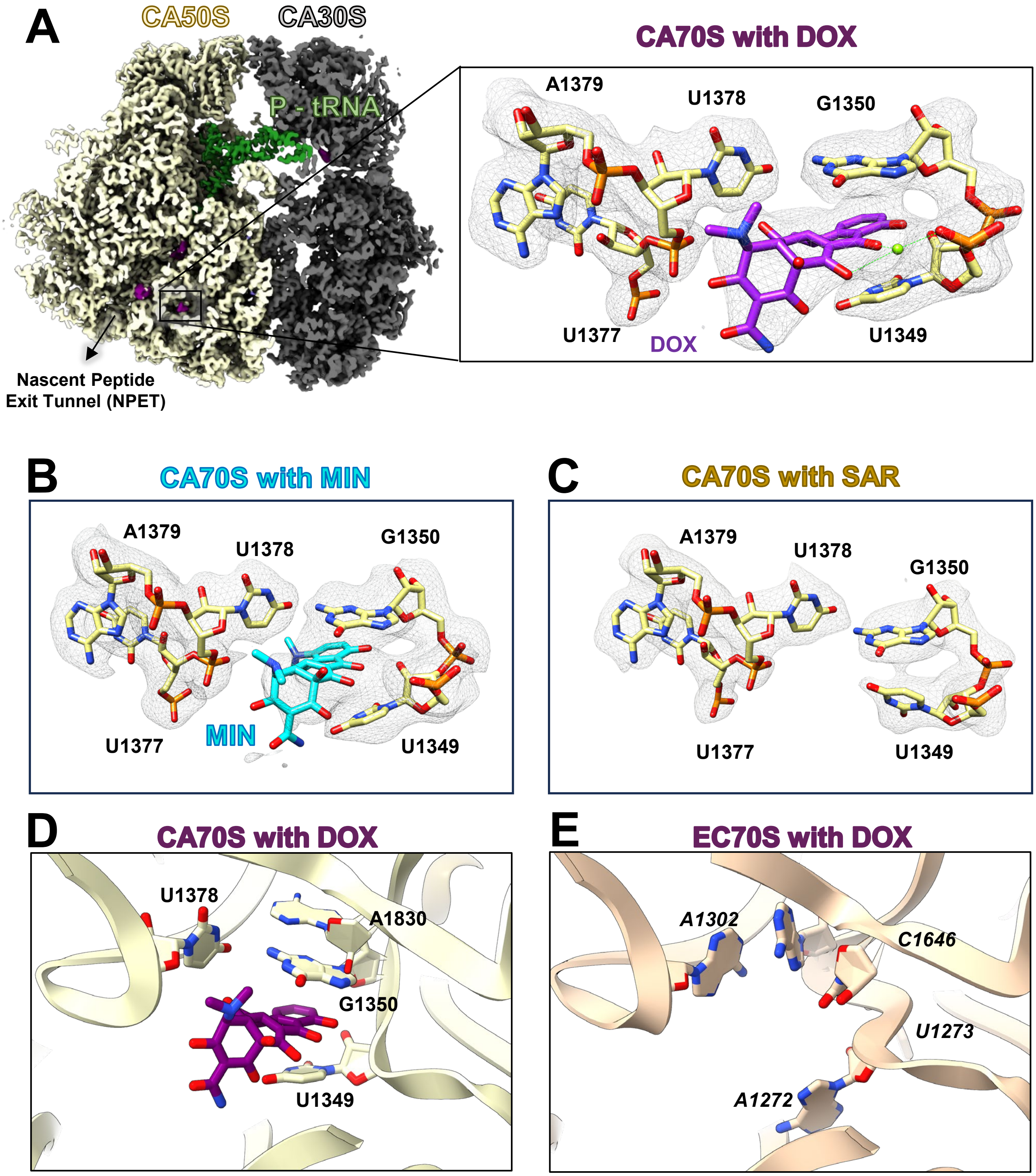

A
CA50S
CA30S
P - tRNA
Nascent Peptide
Exit Tunnel (NPET)
CA70S with DOX
A1379
U1378
G1350
U1377
DOX
U1349
B
CA70S with MIN
A1379
U1378
G1350
U1377
MIN
U1349
C
CA70S with SAR
A1379
U1378
G1350
U1377
U1349
D
CA70S with DOX
U1378
A1830
G1350
U1349
E
EC70S with DOX
A1302
C1646
U1273
A1272

**Figure S4. (A)** Cryo-EM reconstruction of *C. acnes* 70S ribosome bound to Doxycycline is shown. The insets highlight the atomic model, and the corresponding cryo-EM density of a seventh Doxycycline molecule bound by the 23S rRNA region close to the NPET. **(B-C)** A magnified view of the binding site shown in panel A for the *C. acnes* 70S ribosome bound to Minocycline (B) and Sarecycline (C). The structural model is shown in stick format along with the corresponding cryo-EM map density in mesh. Cryo-EM map density corresponding to Minocycline was observed at this site but not for Sarecycline. **(D-E)** A side-by-side comparison of the Doxycycline binding site shown in panel A highlighting the differences between *C. acnes* (D) and *E. coli* 70S ribosome (E). The sequence and conformational differences result in a lack of Doxycycline binding at this site in the *E. coli* 70S ribosome.

### Figure S5

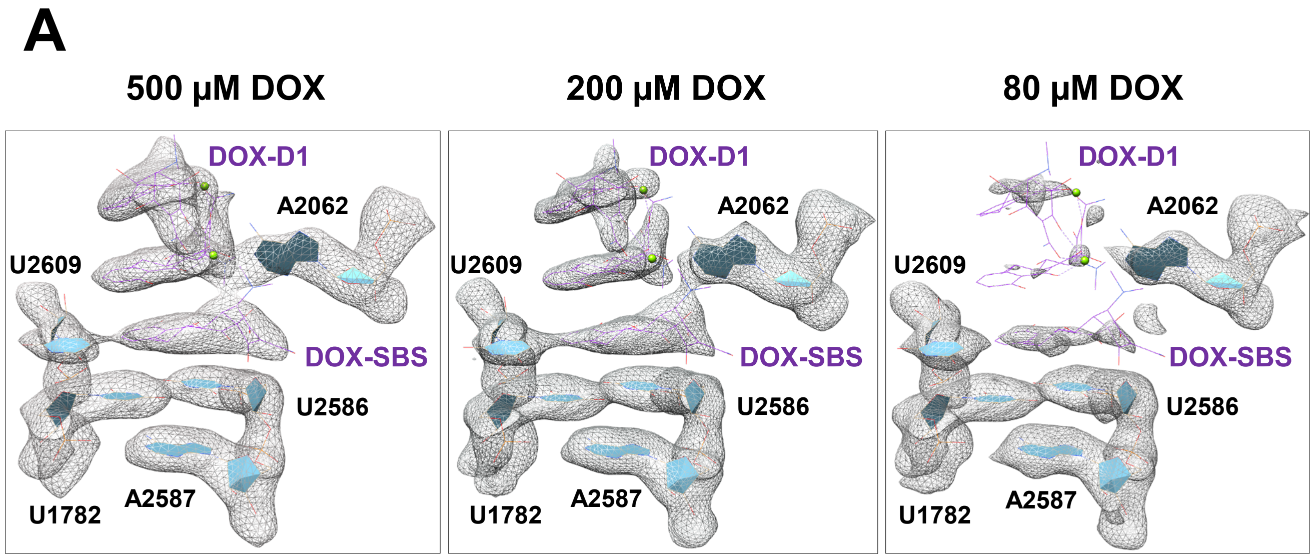


**Figure S5. (A)** A magnified view of the Doxycycline SBS and Dimer 1 binding site for *E. coli* 70S ribosome is shown from three different cryo-EM datasets with varying concentration of Doxycycline (500 µM, 200 µM, and 80 µM). The structural model is depicted in wireframe format and the corresponding cryo-EM map density is shown in mesh.

### Figure S6

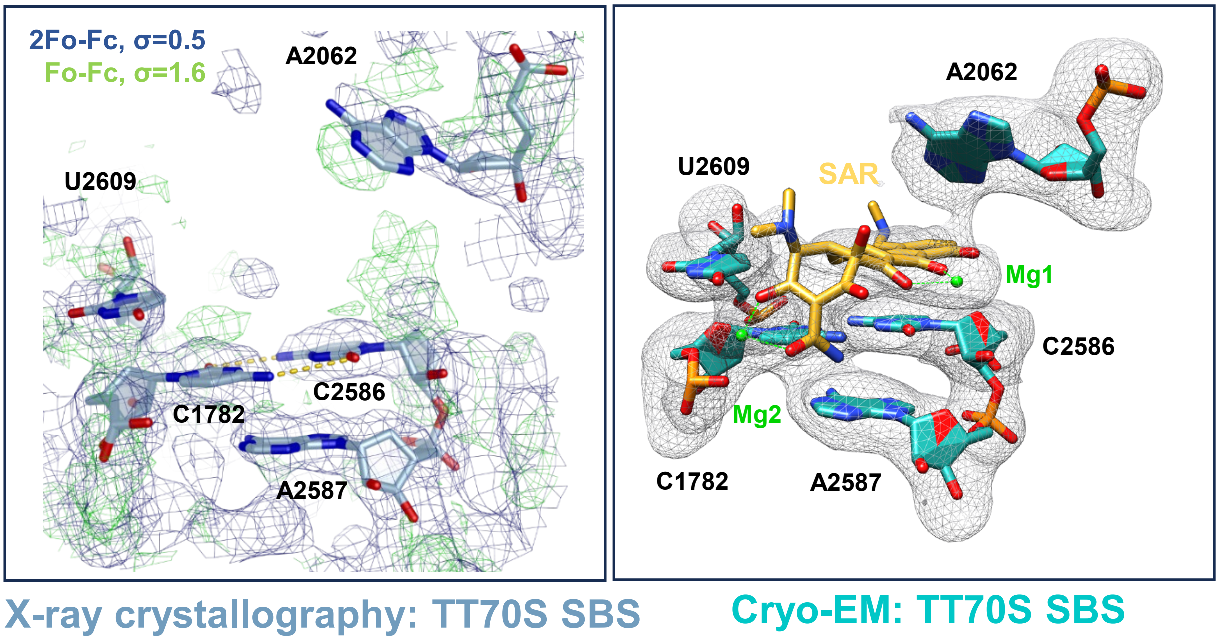


**Figure S6.** A magnified view of the Sarecycline SBS for *T. thermophilus* 70S ribosome is shown from an X-ray crystal structure (left panel) and a cryo-EM structure (right panel). The structural model is depicted in stick format and the corresponding cryo-EM map density is shown in mesh.

### Table S1 Cryo-EM Data Collection and Processing

| | CA70S-Minocycline | CA70S-Sarecycline | CA70S-Doxycycline | EC70S-Minocycline | EC70S-Sarecycline | EC70S-Doxycycline |
|---|---|---|---|---|---|---|
| **Ligands** | Minocycline | Sarecycline | Doxycycline | Minocycline | Sarecycline | Doxycycline |
| **Number of micrographs** | 5284 | 3969 | 3531 | 4938 | 3756 | 4433 |
| **Magnification** | 81,000x | 81,000x | 81,000x | 81,000x | 81,000x | 81,000x |
| **Voltage (kV)** | 300 kV | 300 kV | 300 kV | 300 kV | 300 kV | 300 kV |
| **Electron exposure ($e^-$/Å$^2$)** | 50 $e^-$/Å$^2$ | 50 $e^-$/Å$^2$ | 50 $e^-$/Å$^2$ | 50 $e^-$/Å$^2$ | 50 $e^-$/Å$^2$ | 50 $e^-$/Å$^2$ |
| **Defocus range (μm)** | - 0.8 to -1.8 | - 0.8 to -1.8 | - 0.8 to -1.8 | - 0.8 to -1.8 | - 0.8 to -1.8 | - 0.8 to -1.8 |
| **Pixel size (Å)** | 1.07 Å physical pixel size | 1.07 Å physical pixel size | 1.07 Å physical pixel size | 1.07 Å physical pixel size | 1.07 Å physical pixel size | 1.07 Å physical pixel size |
| **Microscope** | Titan Krios | Titan Krios | Titan Krios | Titan Krios | Titan Krios | Titan Krios |
| **Detector** | Gatan K3 | Gatan K3 | Gatan K3 | Gatan K3 | Gatan K3 | Gatan K3 |
| **Data Collection Software** | EPU | EPU | EPU | EPU | EPU | EPU |
| **Data Processing Software** | cryoSPARC v4 | cryoSPARC v4 | cryoSPARC v4 | cryoSPARC v4 | cryoSPARC v4 | cryoSPARC v4 |
| **Symmetry imposed** | C1 | C1 | C1 | C1 | C1 | C1 |
| **Initial particle images (no.)** | 890,895 | 862,246 | 914,093 | 1,225,779 | 552,873 | 864,223 |
| **Final particle images (no.)** | 105,824 | 80,548 | 103,369 | 628,487 | 234,706 | 248,202 |
| **FSC threshold and Map resolution (Å)** | 0.143 FSC cutoff<br>2.3 Å | 0.143 FSC cutoff<br>2.5 Å | 0.143 FSC cutoff<br>2.4 Å | 0.143 FSC cutoff<br>2.2 Å | 0.143 FSC cutoff<br>2.4 Å | 0.143 FSC cutoff<br>2.4 Å |
| **Resolution range (Å)** | 2.2 Å – 5 Å | 2.2 Å – 5 Å | 2.2 Å – 5 Å | 2.2 Å – 5 Å | 2.2 Å – 5 Å | 2.2 Å – 5 Å |

Table S1. Details of cryo-EM data collection and data processing for the six datasets are presented in a tabular format.

#### Table S2 Model Building Statistics and Accession Codes

| | CA70S-Minocycline | CA70S-Sarecycline | CA70S-Doxycycline | EC70S-Minocycline | EC70S-Sarecycline | EC70S-Doxycycline |
|---|---|---|---|---|---|---|
| **Ligands** | Minocycline | Sarecycline | Doxycycline | Minocycline | Sarecycline | Doxycycline |
| **Deposition ID** | PDB: 9PJ9<br>EMD-71684 | PDB: 9PJ8<br>EMD-71683 | PDB: 9PJ7<br>EMD-71682 | PDB: 9PIJ<br>EMD-71669 | PDB: 9PII<br>EMD-71668 | PDB: 9PIH<br>EMD-71667 |
| **Map sharpening B factor (Å²)** | - 4 | - 6 | - 3 | - 12 | - 4 | - 7 |
| **Initial Model Used** | PDB: 8CRX | PDB: 8CRX | PDB: 8CRX | PDB: 7K00 | PDB: 7K00 | PDB: 7K00 |
| **Model composition** | | | | | | |
| **Non-hydrogen atoms** | 144436 | 144415 | 144683 | 141677 | 141212 | 141462 |
| **Protein residues** | 5746 | 5746 | 5746 | 5587 | 5587 | 5587 |
| **Nucleotides** | 4620 | 4620 | 4620 | 4457 | 4457 | 4457 |
| **Metal Ions** | 330 | 307 | 300 | 495 | 389 | 369 |
| **R.m.s. deviations** | | | | | | |
| **Bond lengths (Å)** | 0.007 | 0.006 | 0.006 | 0.007 | 0.005 | 0.007 |
| **Bond angles (°)** | 0.66 | 0.74 | 0.74 | 0.64 | 0.58 | 0.59 |
| **Validation** | | | | | | |
| **MolProbity score** | 2.12 | 2.39 | 2.39 | 1.87 | 1.92 | 1.86 |
| **Clashscore** | 7.84 | 8.13 | 8.13 | 5.41 | 5.88 | 5.47 |
| **Ramachandran plot** | | | | | | |
| **Favored (%)** | 94.28 | 93.84 | 93.84 | 96.44 | 96.44 | 96.44 |
| **Allowed (%)** | 5.33 | 5.79 | 5.79 | 3.52 | 3.52 | 3.52 |
| **Disallowed (%)** | 0.39 | 0.37 | 0.37 | 0.04 | 0.04 | 0.04 |

Table S2. Model building, data deposition and validation statistics.

### Table S3 Tetracycline CBS and SBS Occupancy Analysis

| ***E. coli*** **70S ribosome** | | | | ***C. acnes*** **70S ribosome** | |
|---|---|---|---|---|---|
| **Tetracycline Concentration** | **500 μM** | **200 μM** | **80 μM** | **Tetracycline Concentration** | **500 μM** |
| **Minocycline CBS** | 1.00 (ref.) | 0.97±0.00 | 0.89±0.02 | **Minocycline CBS** | 0.94±0.01 |
| **Minocycline SBS** | 0.49±0.01 | 0.11±0.05 | 0 | **Minocycline SBS** | 0.84±0.03 |
| **Sarecycline CBS** | 0.85±0.01 | 0.79±0.01 | 0.47±0.11 | **Sarecycline CBS** | 0.88±0.01 |
| **Sarecycline SBS** | 0.34±0.04 | 0.1±0.05 | 0 | **Sarecycline SBS** | 0.30±0.01 |
| **Doxycycline CBS** | 0.97±0.03 | 0.86±0.04 | 0.73±0.05 | **Doxycycline CBS** | 0.96±0.01 |
| **Doxycycline SBS** | 0.60±0.02 | 0.35±0.01 | 0 | **Doxycycline SBS** | 0.54±0.04 |
| **Doxycycline Dimer 1** | 0.43±0.01 | 0.14±0.05 | 0 | **Doxycycline Dimer 1** | 0.28±0.02 |
| **Doxycycline Dimer 2** | 0.26±0.03 | 0.14±0.01 | 0 | **Doxycycline Dimer 2** | 0.34±0.01 |

Table S3. The mean charge density (CD) ratios between antibiotic and neighboring RNA backbone plots were calculated and rescaled to the reference of Minocycline CBS data. The asymptotic ratios as a function of increasing ΔB value of Gaussian smoothing function should represent the relative occupancy of the antibiotic to rRNA nucleotides at a given site.

#### References


1. D. N. Mastronarde, Automated electron microscope tomography using robust prediction of specimen movements. *J Struct Biol* **152**, 36-51 (2005).
2. A. Punjani, J. L. Rubinstein, D. J. Fleet, M. A. Brubaker, cryoSPARC: algorithms for rapid unsupervised cryo-EM structure determination. *Nat Methods* **14**, 290-296 (2017).
3. Z. L. Watson *et al.*, Structure of the bacterial ribosome at 2 A resolution. *Elife* **9**, (2020).
4. I. B. Lomakin, S. C. Devarkar, S. Patel, A. Grada, C. G. Bunick, Sarecycline inhibits protein translation in Cutibacterium acnes 70S ribosome using a two-site mechanism. *Nucleic Acids Res* **51**, 2915-2930 (2023).
5. P. Emsley, K. Cowtan, Coot: model-building tools for molecular graphics. *Acta crystallographica. Section D, Biological crystallography* **60**, 2126-2132 (2004).
6. P. D. Adams *et al.*, PHENIX: a comprehensive Python-based system for macromolecular structure solution. *Acta crystallographica. Section D, Biological crystallography* **66**, 213-221 (2010).
7. E. F. Pettersen *et al.*, UCSF Chimera--a visualization system for exploratory research and analysis. *J Comput Chem* **25**, 1605-1612 (2004).
8. E. F. Pettersen *et al.*, UCSF ChimeraX: Structure visualization for researchers, educators, and developers. *Protein Sci* **30**, 70-82 (2021).
9. M. D. Winn *et al.*, Overview of the CCP4 suite and current developments. *Acta Crystallogr D Biol Crystallogr* **67**, 235-242 (2011).
10. E. F. Pettersen *et al.*, UCSF Chimera - A visualization system for exploratory research and analysis. *J Comput Chem* **25**, 1605-1612 (2004).
11. J. Wang, Experimental charge density from electron microscopic maps. *Protein Sci* **26**, 1619-1626 (2017).
